# Multigenerational effect of heat stress on the *Drosophila melanogaster* sperm proteome

**DOI:** 10.1101/2022.10.20.513068

**Authors:** Shagufta Khan, Rakesh K Mishra

## Abstract

The notion that genes are the sole units of heredity and that a barrier exists between soma and germline has been a major hurdle in elucidating the heritability of traits that were observed to follow a non-Mendelian inheritance pattern. It was only after the conception of epigenetics by Conrad Waddington that the effect of parental environment on subsequent generations via non-DNA sequence-based mechanisms, such as DNA methylation, chromatin modifications, non-coding RNAs and proteins, could be established, now referred to as multigenerational epigenetic inheritance. Despite growing evidence, the male gamete-derived epigenetic factors that mediate the transmission of such phenotypes are seldom explored, particularly in the model organism *Drosophila melanogaster*. Using the heat stress-induced multigenerational epigenetic inheritance paradigm in a widely used position-effect variegation line of *Drosophila*, named *white*-*mottled*, we have dissected the effect of heat stress on the sperm proteome in the current study. We demonstrate that multiple successive generations of heat stress at the early embryonic stage results in a significant downregulation of proteins associated with a diverse set of functions, such as translation, chromatin organization, microtubule-based processes, and generation of metabolites and energy, in the sperms. Based on our findings, we propose chromatin-based epigenetic mechanisms, a well-established mechanism for multigenerational effects, as a plausible way of transmitting heat stress memory via the male germline in this case. Moreover, we show that despite these heat stress-induced changes, the life-history traits, such as reproductive fitness and stress tolerance of the subsequent generations, are unaffected, probing the evolutionary relevance of multigenerational epigenetic effects.

## INTRODUCTION

Organisms frequently encounter stressful environmental conditions such as temperature extremes, toxins, oxidative stress, and nutrient deprivation. Safeguarding mechanisms from damage by such environments are hardwired in an organism’s genome to ensure survival and propagation. This is accomplished by significant gene expression and physiological changes at the cellular level, collectively called the cellular stress response. For instance, the cellular stress response to high temperature encompasses an overall decline in transcription and translation and enhanced degradation of damaged or misfolded proteins. Concurrently, the expression of stress response genes, such as heat shock protein (HSP) coding genes, is elevated (1–3). Usually, after the culmination of stress, cells recuperate by restoring the gene expression status to that of a standard scenario. However, emerging evidence suggests that specific stress-induced changes are not only remembered throughout the lifetime of an organism but are also inherited by subsequent generations, having the potential to be adaptive or maladaptive.

The influence of the parental environment on the offspring via non-DNA sequence-based or epigenetic mechanisms has been shown in various organisms, such as plants (4, 5), nematodes (6, 7), insects (8, 9), and vertebrates (10–13). Such effects, broadly referred to as multigenerational epigenetic effects, are intergenerational or transgenerational based on the timescale of inheritance. They are classified as intergenerational when the effects are only seen in the generations whose genetic material has been directly exposed to the environmental trigger. In contrast, they are transgenerational when the effects are also seen in the stress-free generations whose genetic material has not been directly exposed to the trigger (14). A growing body of literature has deciphered several molecular signals that bring about multigenerational epigenetic inheritance (MEI). They include covalent modifications of DNA, such as adenine and cytosine methylation, and histones, such as methylation, acetylation, phosphorylation, and ubiquitination, and *trans*-acting factors, such as non-coding RNAs (ncRNAs) and proteins (15–17). Moreover, these signals have been suggested to follow either the ‘replicative’ or ‘reconstructive’ mode for the inheritance of multigenerational effects. In ‘replicative’ mode, the epigenetic signal is transmitted through meiosis in its primary mitotic form. In ‘reconstructive’ transmission, the primary epigenetic signals are erased during reprogramming events but are faithfully established in the progeny based on a secondary signal (17, 18). Though our understanding of MEI has revealed several attributes shared between organisms, such as molecular signals and modes of transmission, it has also pointed out the disparity in the MEI mechanisms preferred by different organisms.

*Drosophila melanogaster*, a well-established model organism, is now increasingly used to study multigenerational epigenetic effects owing to its short life cycle, powerful genetic tools, ability to produce many progenies, and evolutionarily conserved epigenetic mechanisms. Despite that, the molecular signals that facilitate the male gamete-mediated inheritance of stress-induced epigenetic effects have seldom been explored in *Drosophila* (19). In the present study, we have employed a previously reported (8) paradigm of heat stress-induced MEI in *Drosophila*. Heat stress-induced MEI in *Drosophila* was demonstrated using position-effect variegation (PEV) line (20, 21), referred to as *white-mottled* (22, 23). In this X-chromosomal inversion line, the organisms exhibit a mottled eye phenotype (red and white patches) caused by the juxtaposition of the *white* gene near pericentromeric heterochromatin. Although previous studies have revealed that heat stress at early embryonic stages suppresses PEV in *Drosophila* (24, 25), the underlying mechanism was revealed later by Seong *et al*. (8). They demonstrated by measuring the eye pigment levels, which serves as a reliable proxy for the expression state of the *white* gene, that heat stress at 0-3 hour embryonic stage disrupts the structure of heterochromatin via Mekk1-p38 mediated phosphorylation of the activating transcription factor-2, required for the formation of heterochromatin together with heterochromatin protein 1. Interestingly, these epigenetic changes were not only maintained till adulthood, surpassing the mitotic divisions that happen during development, but were also shown to get inherited by subsequent generations via the male germline. However, an insight into the signals that bring about heat stress-induced MEI via the male germline of *Drosophila* is still lacking.

Therefore, we first re-examined and confirmed the transmission of heat stress-induced MEI via the male germline by assaying the eye pigment levels of the males subjected to multigenerational heat stress (**Figure 1A**). Next, deciphered the underlying molecular signals in the male germline of *Drosophila*. To this end, using liquid chromatography with tandem mass spectrometry (LC-MS/MS) analysis, we identified heat stress-induced changes in the sperm proteome of the *white-mottled* strain. Our data shows substantial downregulation of proteins belonging to a diverse set of functions in the sperms in response to multiple generations of heat stress. Furthermore, we shed light on the adaptive or maladaptive potential of heat stress-induced epigenetic effect by assessing the reproductive potential of the impacted individuals and the heat stress tolerance of their progenies. Overall, our study provides a global view of the sperm proteome dynamics in response to heat stress, presents a way forward in characterizing the sperm-derived factors involved in MEI, and opens new avenues in understanding stress-induced epigenetic inheritance in *D. melanogaster*.

**Figure 1.**
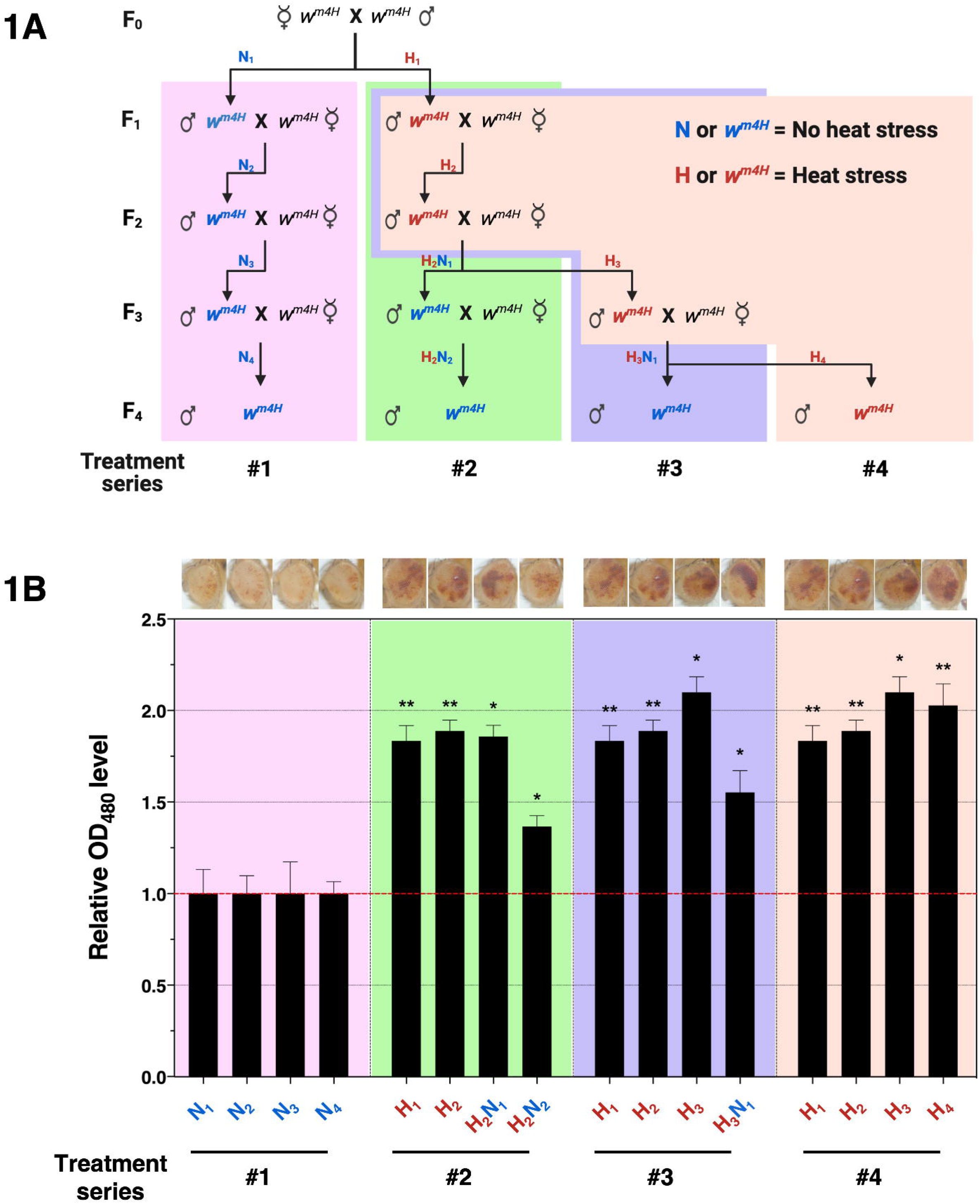
Multigenerational inheritance of heat stress-induced disrupted chromatin state. **(A)** Male and female *w^m4H^* flies (F_0_) were mated to collect 0-3 hour F_1_ embryos. In treatment series #2, the F_1_ embryos were exposed to heat stress (1 hour at 37°C) to obtain H_1_ adults. H_1_ males were then mated with untreated *w^m4H^* females to collect F_2_ embryos (0-3 hour), followed again by an exposure to heat stress to obtain H_2_ adults. Similar matings were repeated until the F_4_ generation without heat stress. In treatment series #3, F_1_, F_2_, and F_3_ embryos were exposed to heat stress whereas, in series #4, embryos were exposed to heat stress at every generation. Unstressed embryos were taken in every generation in treatment series #1 (control). **(B)** In all the treatment series, the amount of eye pigment of 4-days old male progenies was measured and the average eye pigment levels relative to the control males (relative OD_480_) are presented in the bar graph (n = 3). Error bars represent standard error. **p < 0.01; *p < 0.05. Representative PEV phenotypes of the eye are shown above.

## EXPERIMENTAL PROCEDURES

### Drosophila strain and culture

The *Drosophila melanogaster* or fruit fly strains used in this study were *In(1)w^m4H^* (*w^m4H^* for short) (23) and ProtamineA-eGFP (26), kindly provided by Francois Karch (Department of Genetics and Evolution, University of Geneva, Switzerland) and Christina Rathke (Department of Biology, Philipps University of Marburg, Germany), respectively. Fruit flies were cultured on a standard medium containing corn flour, sugar, yeast, malt, agar, and preservatives. The stocks were maintained at a constant density in the food bottles. However, embryos were collected, given heat stress, and allowed to develop in the food vials for heat stress experiments. All the experiment vials were kept at 25°C except during the heat stress window. A 12-hour light-dark cycle was maintained throughout.

### Multigenerational heat stress regime

Crosses were set using twenty virgin females and five males of *w^m4H^* (F_0_). F_1_ embryos were collected for three hours at 25°C in the food vials. After collection, the vials containing the embryos (0-3 hour) were kept for 1 hour inside a 37°C incubator for heat stress. Post-stress, all the vials were maintained at 25°C for further development into adult flies. For unstressed control, sibling embryos (0-3 hour) were collected and allowed to develop into adults at 25°C without heat stress. The stressed (H_1_) and unstressed (N_1_) male progenies thus obtained were mated again with unstressed virgin females in a similar ratio. An identical method of embryo collection and heat stress was performed for four generations based on the mating scheme and heat stress strategy shown in **Figure 1A**.

### Eye pigment quantification and imaging

As previously described (9), four-day old male flies were frozen in liquid nitrogen and subsequently vortexed for decapitation. Heads were manually separated, counted, and transferred into fresh 1.5 ml centrifuge tubes. Next, they were homogenized in 200 μl of 30% ethanol (acidified to pH 2.0) using a micro pestle attached to the hand-held homogenizer. After one hour of incubation on a rotator at room temperature (RT) in the dark, the tubes were centrifuged twice for 3 minutes at 12,000g, transferring the supernatant into a fresh tube each time. The final supernatant was used to measure the absorbance at 480 nm (OD_480_). Acidified 30% ethanol (pH 2.0) was used as blank. The absolute OD_480_ values were divided by the number of heads in the corresponding tube and then used for further analysis. We used the student’s t-test with Welch’s correction to calculate the statistical significance. For statistical analysis and generating graphs, GraphPad Prism 7 software was used. The four-day old male flies were frozen in liquid nitrogen, and their eyes were imaged using Axio Zoom.V16 microscope (Zeiss).

### Isolation of sperms

Mature sperms from the seminal vesicles of *w^m4H^* male flies were isolated as previously described (27, 28). In brief, male *w^m4H^* flies, kept in separate food vials for 5-10 days post eclosion, were anesthetized on a CO_2_ pad of the stereomicroscope. In two cavity blocks, pre-rinsed once with 70% ethanol and twice with Milli-Q, 3 ml of ice-cold 1X PBS containing 1mM PMSF was taken. One cavity block was kept on ice to collect the seminal vesicles, and another was kept under the microscope. Males were gently picked with the forceps between the thorax and abdomen (Mini Dumont #M5S) and dipped into the cavity block under the microscope. An incision was made between the last two abdominal segments using an insulin needle. The male reproductive tract, consisting of the testis, seminal vesicles, and accessory glands, was pulled out with the needle. After removing the accompanying tissues, such as accessory glands, the seminal vesicles were separated from the testis and transferred into the cavity block kept on ice. Once the seminal vesicles from 50 flies were collected (~40 minutes), they were punctured with a needle. The cloudy mass of sperms was pulled away from the epithelial layer using the needle and transferred into a 1.5 ml microfuge tube kept on the ice having 100 μl of 1X PBS containing 1X protease inhibitor cocktail (PIC) (cOmplete EDTA-free, Merck). The sperms isolated from 75-100 male flies were used for further processing.

### Squash preparation of testes

Testes squashes of the ProtamineA-eGFP line of *Drosophila* were prepared as previously described (29), with minor modifications. Briefly, testes were dissected in 1X PBS and fixed for 30 minutes in 4% formaldehyde. Next, they were transferred onto a coverslip in a drop of 1X PBS and squashed by placing a slide on the top. The slides were frozen in liquid nitrogen, after which the cover slip was removed. The slides were incubated in Coplin jars at −20°C in methanol for 5 minutes, followed by 1/1 (v/v) methanol/acetone for 5 minutes, and finally into acetone for 5 minutes. The slides were washed once with 1X PBS containing 0.1% Triton-X 100 at RT for 10 minutes and then twice with 1X PBS at RT for 5 minutes. The slides were stained with DAPI and imaged using the Leica TCS SP8 confocal microscope.

### Protein extraction and estimation

The isolated sperms were centrifuged for 5 minutes at 12,000 rpm at 4°C, and the supernatant was discarded. The sperm pellet was washed thrice with ice-cold 1X PBS containing 1X PIC, discarding the supernatant each time after centrifugation. The final sperm pellet was re-suspended in 120 μl of lysis buffer (10 mM Tris-Cl pH 8.0, 1 mM EDTA pH 8.0, 10 mM NaCl, 1% SDS and 40 mM DTT) and incubated at 37°C until clear (~1 hour). Next, Laemmli buffer was added to the permeabilized sperms and incubated at 95°C for 5 minutes. The whole sperm protein extract thus obtained was quantified using Pierce™ 660nm protein assay reagent (Thermo Scientific™) according to the manufacturer’s instructions.

### Western blot analysis

Whole sperm protein extracts were resolved on 12% SDS-PAGE gel, according to the standard protocol (30). The proteins were transferred from polyacrylamide gel onto a polyvinylidene difluoride membrane (Millipore IPVH00010). After the transfer, the membranes were stained with Anti-Lamin Dm0 (DSHB, ADL67.10) and Anti-ß-tubulin (DSHB, E7) antibodies. In brief, the membranes were incubated in 5% Blotto made in 1X Tris-buffered saline with 0.1% Tween 20 detergent (TBST) for blocking at RT for 1 hour. Primary antibody was added at the desired concentration (1:1000 for Anti-Lamin Dm0 and 1:400 for Anti-ß-tubulin) in 1X TBST and incubated at 4°C with the membrane overnight. The next day, membranes were washed thrice for 7 mins each with 1X TBST at RT. The Anti-mouse HRP secondary antibody (Abcam, ab97046) was added at 1:10,000 concentration in 0.5% Blotto made in 1X TBST and incubated with the membrane for 1 hour at RT. The membrane was rewashed three times for 7 mins each with 1X TBST. The blot was developed using the ECL Prime reagent (GE healthcare - RPN2236), and the image was captured in chemicapt. The images were processed using Adobe Photoshop.

### In-gel digestion and identification of proteins by LC-MS/MS

The samples for LC-MS/MS analysis were prepared and run as previously described (31, 32), with minor changes. Briefly, 30 μg of whole sperm protein extract was resolved on 12% SDS-PAGE gel, following the standard protocol (30). After staining the gel with Imperial™ Protein Stain (Thermo Scientific), each lane was sliced into four pieces giving rise to four fractions per sample. The gel pieces were cut into cubes of ~1 mm^3^ size and were destained and dehydrated using 50% acetonitrile (ACN) and 100% ACN, respectively. The proteins in the destained and dehydrated gel pieces were reduced by adding 10 mM DTT, followed by alkylation using 55 mM iodoacetamide. The gel pieces were again dehydrated using 50% ACN, followed by 100% ACN. Next, protein in the gel pieces was digested using 20 ng/μl of Trypsin solution made in 25mM ammonium bicarbonate (Promega, Madison, WI) by incubating at 37°C overnight. The digested peptides were extracted in 50% ACN and 2.5% TFA, vacuum dried, and resuspended in 100 μl of 0.1% TFA and 2% ACN. Resuspended peptides were desalted using 100 μl Pierce™ C18 pipette tips (Thermo Scientific™) following the manufacturer’s instructions. The peptides were then vacuum dried and resuspended in 20 μl of 2% ACN and 0.1% formic acid for loading onto the Q-Exactive™ HF Mass Spectrometer (Thermo Scientific™).

Trypsin-digested peptides were loaded on PepMap™ RSLC C18 columns (3 μm, 100Å, 75 μm X 15cm). 0 to 90% acetonitrile gradient of 90 min length was used to separate the peptides. The Easy-nLC 1200™ system (Thermo Scientific) was connected with a Thermo Scientific Q-Exactive™ HF instrument via a nano-electrospray source, operated at 2.2 kV. The ion transfer tube was operated at 300°C. The data-dependent mode was selected to run the mass spectrometer. The MS2 scans were obtained with a resolution of 15,000 at m/z 200−2000 in an Orbitrap mass analyzer. In an acquisition cycle, ten precursors containing double or higher charge states were chosen for sequencing and fragmented in the Orbitrap using higher energy collisional dissociation with 28% normalized collision energy. To decrease the possibilities of repeated sequencing, dynamic exclusion was activated for all sequencing events. Peaks that were selected for fragmentation more than once within 15 seconds were excluded.

### Analysis of LC-MS/MS data

The proteome profile was recorded in biological triplicates. MaxQuant (version 2.0.3.0) (33, 34) with an integrated andromeda peptide search engine (35) was used to process the raw spectra. *D. melanogaster* reference proteome, downloaded from UniProtKB (Proteome ID – UP000000803), containing 23,524 entries was used to perform the search. Trypsin/P was selected as the cleavage enzyme, with a maximum of two missed cleavages. Cysteine carbamidomethyl was set as the fixed modification, and oxidation of methionine and N-terminal acetylation as the variable modifications. False discovery rate (FDR), initial precursor mass tolerance, and fragment mass deviation were set to 1%, 20 ppm, and 0.5 Da, respectively. All other parameters were set to default in MaxQuant. Label-free quantification (LFQ) option was selected for relative quantification based on peptide intensities.

### Experiment design and statistical rationale

Biological triplicates (decided based on sample handling, quality, and minimum requirement for statistical analysis) from independent experiments were analyzed for each treatment group of F_1_ (N_1_ and H_1_) and F_4_ (N_4_, H_2_N_2_, H_3_N_1_, and H_4_) generation. Since the LC-MS/MS experiments were conducted at different time points, the comparative analysis was done separately between the treated and untreated groups of the F_1_ and F_4_ generations. The output ‘proteingroups.txt’ files generated by MaxQuant were used for further analysis with Perseus software (version 1.6.15.0) (36, 37). The proteins were first filtered to remove the ‘potential contaminants’, ‘only identified by site’, and ‘reverse’ positive hits. Distribution plots were generated using the log2 transformed LFQ intensity values. Pearson’s correlation coefficient was computed (since the data followed a normal distribution) to assess the reliability between the biological triplicates. For unique peptide-based comparative analysis between the treatment groups, proteins identified with two or more unique peptides were considered. For LFQ-based analysis between the groups, the log2 transformed LFQ intensity values were further filtered using Perseus software for a minimum of one valid value in at least one treatment group. Following this, median-based column and row normalization (Z-score) were done within and across treatment groups, respectively. Two-sample student’s t-tests were performed between the treatment groups, using the default parameters (S0 = 0, Permutation-based FDR = 0.05) to obtain the significant values. The significantly different proteins were used to plot a heat map using default parameters of hierarchical clustering in Perseus.

### Gene ontology analysis

DAVID (version 2021), a web-based functional annotation tool, was used to do the gene ontology (GO) enrichment analysis (38, 39). Flybase gene identifiers were uploaded to DAVID, and the default *D. melanogaster* gene list was selected as the background. The analysis was performed using the Biological Process (BP) as the annotation category and default run parameters. The annotation terms ≤ 0.01 FDR and Benjamini-Hochberg corrected p-value < 0.01 were further considered for biological interpretation.

### Measurement of reproductive fitness and heat stress tolerance

Three *w^m4H^* males (two-day old) from each treatment group in every generation (F_1_ to F_4_), as represented in **Figure 1A**, were mated with ten age-matched virgin *w^m4H^* females (n = 5). Embryos were collected for three hours at 25°C in the food vials. After collection, the embryos were counted under a stereomicroscope to determine fecundity. The embryos were kept at 25°C for further development into adults. To determine fertility, the number of adults emerging from each vial was counted and divided by the number of embryos in the corresponding vial. For assessing heat stress tolerance, sibling embryos were collected in a similar manner and subjected to heat stress for 1 hour at 37°C. After exposure, vials were kept at 25°C for further development. The number of adults from each vial was counted and divided by the number of embryos in the corresponding vial. To calculate the statistical significance, student’s t-test with Welch’s correction was used. For statistical analysis and generating graphs, GraphPad Prism 7 software was used.

## RESULTS

### Heat stress-induced multigenerational epigenetic inheritance

Based on the previous study reporting the multigenerational effect of heat stress in the *w^m4H^* line of *D. melanogaster* (8), we subjected the *w^m4H^* embryos at an early stage (0-3 hour) to multigenerational heat stress as presented in **Figure 1A**. Since two generations of heat stress were shown to result in a transgenerational effect, we designed the experiments where the embryos were subjected to stress for four generations (treatment series #4), three generations followed by one stress-free generation (treatment series #3), two generations followed by two stress-free generations (treatment series #2), and no heat stress for four generations (treatment series #1). Consistent with the previous report (8), we observed that exposing F_1_ embryos to heat stress results in a significant increase in the eye pigment levels of the adult males (H_1_) when compared to the unexposed male siblings (N_1_) (**Figure 1B**). A similar increase in eye pigment levels was observed upon exposure to heat stress in every generation for four generations (treatment series #4, H_1_ to H_4_). Moreover, the eye pigment levels remained significantly higher in the progenies of stressed males even in the absence of stress for one generation (treatment series #3, H_1_ to H_3_N_1_) or two generations (treatment series #2, H_1_ to H_2_N_2_) when compared to the corresponding generation’s unstressed males (treatment series #1, N_1_ to N_4_). These results indicate and corroborate with the previous findings that heat stress at an early embryonic stage results in the desilencing of the *white* gene in *w^m4h^* flies, as shown by the eye pigment assay, and that this epigenetic change (albeit not as an expression state of the *white* locus, see **Figure S1**) is multigenerationally inherited to subsequent generations via the male germline or sperm.

The MEI of heat stress-induced changes via the male germline in the *w^m4H^* flies, hereafter referred to as heat stress memory for simplicity, represents a reconstructive mode of transmission of epigenetic information (17, 18). This is because the unstressed male progenies of heat-stressed males exhibit high eye pigment levels regardless of inheriting the X-chromosome, which carries the *white* locus under scrutiny, from an unstressed female parent (**Figure S1A**), possibly via *trans*-communication between the stressed autosomal chromosomes and unstressed X-chromosome (**Figure S1B**). Therefore, to identify the potential secondary signals in the milieu of sperms that communicate the heat stress memory to the subsequent generations and thereby alter the expression status of the *white* gene, we next isolated the sperms of F_1_ and F_4_ generations from all the treatment series, assessed the purity of the sperm isolates and performed LC-MS/MS analysis.

### Quality assessment of the sperm isolates

The sperms were isolated from the seminal vesicles of stressed and unstressed *w^m4H^* males from the F_1_ and F_4_ generations (**Figure 2A**). The purity of the sperm isolates was assessed at two stages of sample preparation. The isolates were first checked for somatic cell contamination and cells from the early stages of spermatogenesis by looking at the nuclear morphology of the dissected sperm mass from each batch. Since the mature sperms of *D. melanogaster* have distinct needle-shaped nuclei (**Figure 2B**) (26, 40), the absence of nuclear morphology other than the needle shape (**Figure 2C**) indicated the purity of sperm isolates. Secondly, the mature sperms of *D. melanogaster* are devoid of nuclear Lamin Dm0 or Lamin B (41, 42), a major component of the nuclear membrane in the somatic cells and early stages of spermatogenesis. Therefore, the whole sperm protein extracts were checked for the Lamin Dm0 signal. The blots probed against the anti-Lamin Dm0 antibody (**Figure 2D** and **Figure S2**) showed a strong signal in 0-16 hour embryo extracts (positive control for anti-Lamin Dm0 antibody). In contrast, no signal was observed in the whole sperm extracts. Anti-ß-tubulin antibody was used as a loading control for sperm extracts. Together, these results confirm the purity of sperm isolates used in this study for LC-MS/MS analysis.

**Figure 2.**
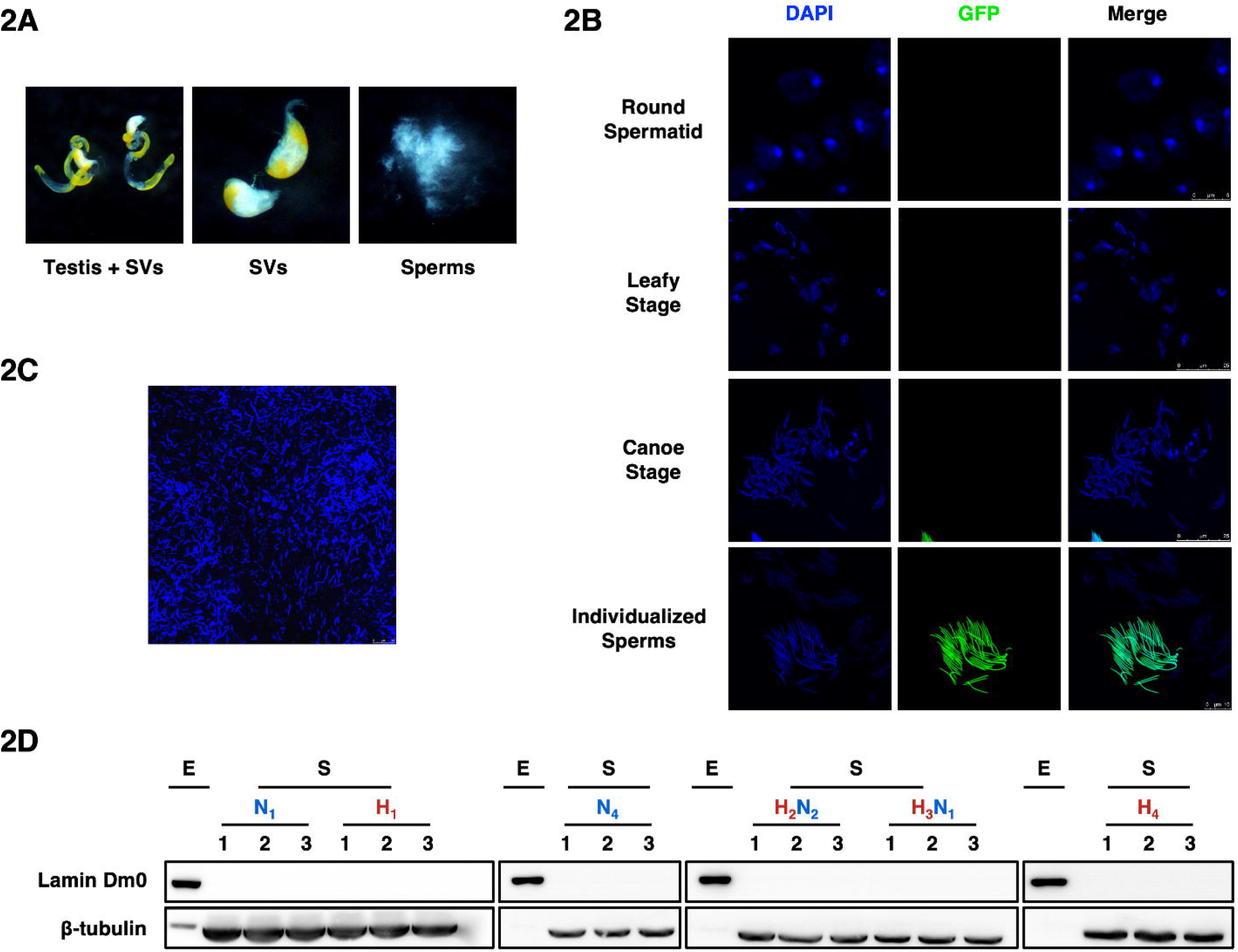
Isolation and quality assessment of sperms. **(A)** Stereomicroscopic images of the different steps of isolation of sperms from the *w^m4H^* male flies. Adults were dissected, and testis plus seminal vesicles (or SVs) were pulled out (left). SVs were separated from the testis (middle) and punctured to release the cloudy mass of sperms (right). **(B)** Changes in the nuclear morphology during spermatogenesis in the testis of the ProtamineA-eGFP transgenic line. Since the protamines are exclusively translated in the individualized sperms, eGFP signal is observed only during the individualization stage. The scale bar is 5 μm for the round spermatid stage, 25 μm for the leafy and canoe stage, and 10 μm for individualized sperms. **(C)** DAPI stained isolated mass of sperms from SVs of *w^m4H^* males indicates that the sperm isolates are devoid of contamination from somatic cells and early stages of spermatogenesis. The scale bar is 25 μm. **(D)** Western blot analysis of the whole sperm protein extracts (S) using anti-Lamin Dm0 and anti-ß-tubulin antibody shows the absence of Lamin Dm0 signal in the sperm extracts, indicating the purity of the isolates. Lamin Dm0 signal in embryo extract (E) is used as a positive control for anti-Lamin Dm0 antibody, whereas anti-ß-tubulin is used as a loading control for sperm extracts.

### Overview of the LC-MS/MS analysis

The unique morphology of the *Drosophila* sperm limited the analysis of sperm nuclei, which constitutes less than 1% (~10 μm) of the whole sperm size (~1.8 mm) (43–45) and is likely the vehicle for factors that act in *trans* on the *white* locus in subsequent generations. Therefore, in the current study, we performed the LC-MS/MS analysis using the whole sperms isolated from the males of all the treatment groups from F_1_ (N_1_ and H_1_) and F_4_ (N_4_, H_2_N_2_, H_3_N_1_, and H_4_) generation. Although this strategy yields a global picture of proteomic changes, we reasoned that it would still unveil general heat stress-induced features. To do this, the output files from MaxQuant of F_1_ and F_4_ generation were filtered to remove ‘potential contaminants’, ‘only identified by site’, and ‘reverse’ hits (**Table S1**). A total of 987, 982, 900, 867, 720, and 741 proteins were identified with at least one peptide across the biological triplicates of N_1_, H_1_, N_4_, H_2_N_2_, H_3_N_1_, and H_4_, respectively (**Table 1, Table S2**, and **Figure S3**). Out of these, 863, 850, 765, 735, 546, and 557 proteins were identified by two or more unique peptides across the biological triplicates of N_1_, H_1_, N_4_, H_2_N_2_, H_3_N_1_, and H_4_, respectively. Relative protein abundances of 807, 800, 678, 681, 524, and 538 proteins were measured by LFQ in N_1_, H_1_, N_4_, H_2_N_2_, H_3_N_1_, and H_4_, respectively.

**Table 1:** Summary of LC-MS/MS profiling.

| Treatment | No. of proteins identified (with two or more unique peptides) |  |  |  |
| --- | --- | --- | --- | --- |
|  | 1 | 2 | 3 | Union |
| **N<sub>1</sub>** | 956 (817) | 927 (783) | 912 (760) | 987 (863) |
| **H<sub>1</sub>** | 929 (781) | 920 (777) | 900 (748) | 982 (850) |
| **N<sub>4</sub>** | 796 (609) | 821 (655) | 846 (832) | 900 (765) |
| **H<sub>2</sub>N<sub>2</sub>** | 813 (803) | 806 (787) | 802 (792) | 867 (735) |
| **H<sub>3</sub>N<sub>1</sub>** | 620 (478) | 660 (494) | 608 (462) | 720 (546) |
| **H<sub>4</sub>** | 647 (474) | 630 (465) | 694 (519) | 741 (557) |

Next, we assessed the quality of the acquired proteomic datasets by checking the distribution of LFQ intensity values in each replicate and computing the correlation coefficients between the replicates of each treatment group. The protein abundances followed a normal distribution (**Figure S4**) and were found to be strongly correlated between the biological triplicates (Pearson’s correlation = 0.94 – 0.99) (**Figure S5**). To further assess the reliability of our data, we compared the unstressed (N_1_ and N_4_) sperm proteomes of *w^m4H^* with the recently reported *D. melanogaster* sperm proteome 3 (DmSP3) (27, 28, 46). The comparison was carried out using the total proteins identified across the biological triplicates of N_1_ and N_4_. Around 96% of the proteins identified in the N_1_ and N_4_ sperm proteome were also identified in DmSP3 (**Figure S6A** and **Table S3**). Additionally, the GO analysis revealed the annotation categories enriched in N_1_ and N_4_ to be a subset of DmSP3 (**Figure S6B** and **Table S3**), further supporting the reliability of our data.

### Effect of multigenerational heat stress on the sperm proteome

To identify the heat stress-induced multigenerational changes in the *w^m4H^* sperm protein composition, we compared the sperm proteomes of stressed and unstressed treatment groups from F_1_ and F_4_ generation using the proteins identified with high confidence (with two or more unique peptides) across the biological triplicates. In addition, we employed an LFQ-based methodology and compared the protein abundances between the stressed and unstressed treatment groups to identify quantitative changes in the sperm proteome. First, we compared the sperm proteome of the individuals that were heat-stressed for one generation at the embryonic stage (H_1_) with that of unstressed individuals of the corresponding generation (N_1_) (**Figure 1A**). Out of the 863 and 850 proteins identified in the F_1_ generation in N_1_ and H_1_, respectively, 814 proteins are common, while 49 are unique to N_1_, and 36 are unique to H_1_ (**Figure 3A** and **Table S4**). Further, we took the sperm proteomes of individuals of the F_4_ generation (H_2_N_2_, H_3_N_1_, and H_4_) that were the carriers of transgenerational heat-stress memory and performed a pairwise comparison with the sperm proteome of unstressed individuals of the corresponding generation (N_4_) (**Figure 1A**). Out of the 765 and 735 proteins identified with high confidence in N_4_ and H_2_N_2_, respectively, 694 proteins are common, while 71 are unique to N_4_ and 41 are unique to H_2_N_2_ (**Figure 3A** and **Table S5**). Similarly, out of the 765 and 546 proteins identified with high confidence in N_4_ and H_3_N_1_, respectively, 531 proteins are common, while 234 and 15 proteins are unique to N_4_ and H_3_N_1_, respectively (**Figure 3A** and **Table S6**). Of the 765 and 557 proteins identified in N_4_ and H_4_, 536 are shared between the two, while 229 and 21 are unique to N_4_ and H_4_, respectively (**Figure 3A** and **Table S7**).

**Figure 3.**
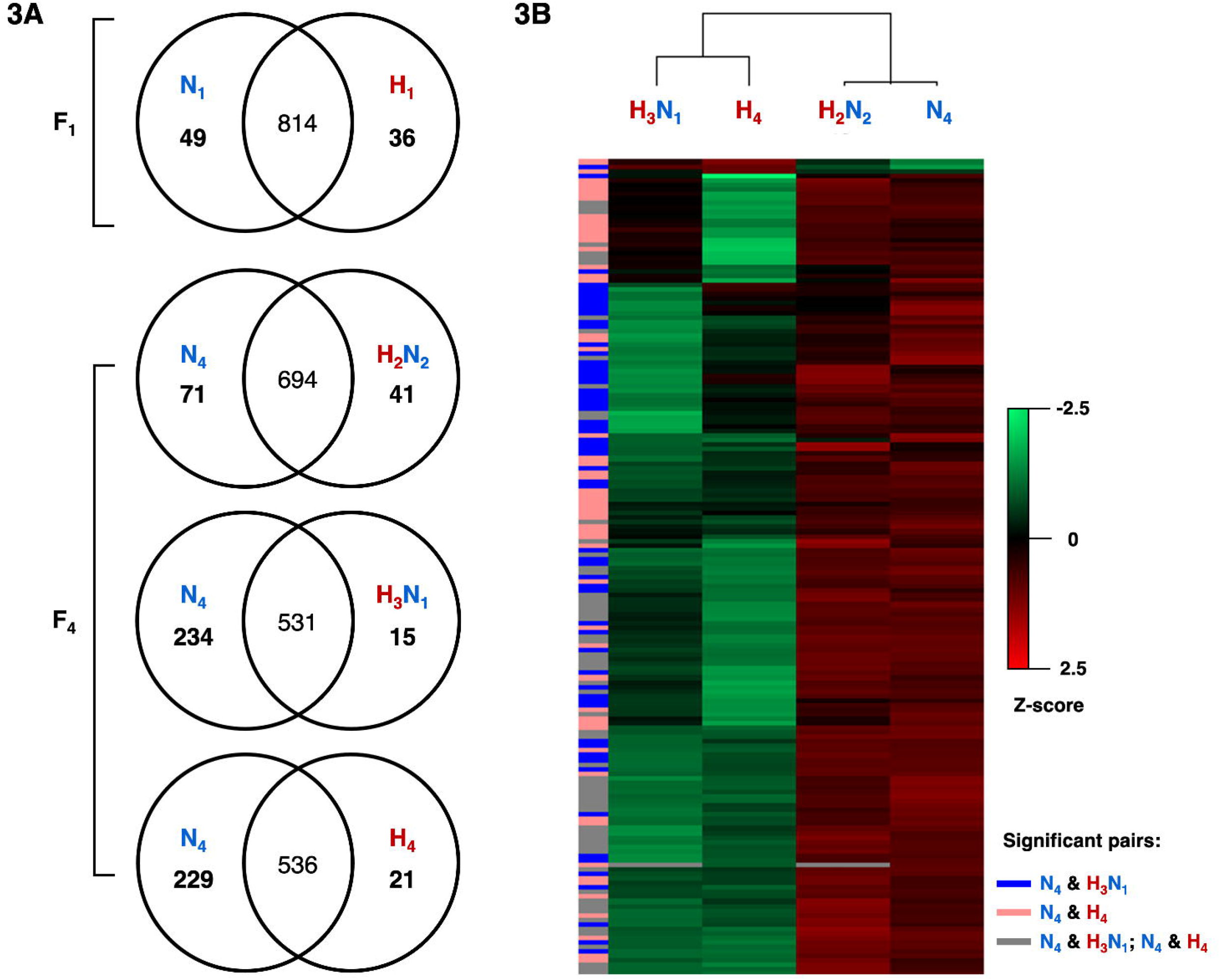
Comparative analysis of sperm proteomes in response to multigenerational heat stress. **(A)** Venn diagrams showing the comparison of *w^m4H^* sperm proteome of unstressed F_1_ males (N_1_) with the stressed F_1_ males that were exposed to heat stress at embryonic stage for one generation (H_1_), and unstressed F_4_ males (N_4_) with the stressed F_4_ males that were exposed to heat stress for two generations followed by two stress-free generations (H_2_N_2_), three generations followed by one stress-free generation (H_3_N_1_), and for four generations (H_4_). **(B)** Heat map representing the relative abundance of proteins that are significantly different in the stressed sperm proteomes of the F_4_ generation (H_2_N_2_, H_3_N_1_, and H_4_) when compared with the unstressed proteome (N_4_). Normalized abundance values of the proteins are represented by red (high abundance) and green (low abundance) color, as indicated in the color scale bar. The missing values are represented in grey color. Significant pairs are indicated on left column. Data represents the median of biological triplicates.

Second, we employed an LFQ-based methodology to investigate the changes in protein abundance in the sperms upon multigenerational heat stress. Amongst the 1007 and 916 proteins identified in the proteome of F_1_ and F_4_ generation (**Table S1**), respectively, we retained 818 and 713 proteins (**Table S8**) after the application of LFQ (see *Experimental Procedures* for criteria). We performed a pairwise student’s t-test using the default parameters to identify the significantly different proteins. In the F_1_ generation, we only found two proteins that were significant and higher in abundance (or upregulated) in H_1_ when compared with N_1_ (**Table S8**). However, in the F_4_ generation, 179 proteins differed significantly in their abundance in response to heat stress. Out of these 179 proteins, 123 and 118 were differentially modulated in H_3_N_1_ and H_4_, respectively, compared with N_4_, with only 62 differentially modulated in both. The hierarchical clustering of these 179 proteins revealed that most proteins were significantly less in abundance (or downregulated) in either H_3_N_1_ or H_4_, or both (**Figure 3B**, **Figure S7**, and **Table S8**). Collectively, our data suggest that multigenerational heat stress changes the protein composition of the sperms, and that the extent of change is proportional to the number of generations exposed to heat stress.

### Functional analysis of differentially regulated proteins

We performed the GO enrichment analysis to identify the biological function associated with the proteins differentially regulated in the F_1_ and F_4_ generation in response to multigenerational heat stress. To do this, we used the upregulated and downregulated proteins in H_1_, H_2_N_2_, H_3_N_1_, and H_4_ – unique peptide-based and LFQ-based comparative analyses were combined, and duplicate protein IDs were removed (**Table S4-S8**). Functional analysis revealed that proteins downregulated in H_3_N_1_ and H_4_ were enriched in a wide variety of BP terms – cytoplasmic translation (small and large ribosomal subunits), DNA-templated transcription initiation, chromatin organization (mainly the core components of nucleosome, and proteins involved in chromatin remodeling during spermatogenesis), microtubule-based process (components of dynein complex, protein required for sperm motility), and generation of energy and metabolites (glycolytic process, electron transport chain, tricarboxylic acid cycle, proton motive force-driven ATP synthesis, and acetyl-CoA biosynthetic process) (**Figure 4** and **Table S9**). Amongst these, only a subset of proteins associated with cytoplasmic translation were also downregulated in H_2_N_2_. Additionally, the BP term sarcomere organization (actin filament and unfolded protein binding) was specifically associated with the proteins downregulated in H_2_N_2_. Proteins involved in multicellular organism reproduction, such as C-type lectin-like proteins, accessory gland proteins, and serpins, were specifically downregulated in H_1_, with a subset of them also downregulated in H_4_. On the other hand, for the proteins upregulated in H_1_, H_2_N_2_, H_3_N_1_, and H_4_, no enriched GOBP categories could be identified. The possible implications of differentially regulated proteins, mainly the ones that are commonly deregulated in H_3_N_1_, and H_4_, are discussed further in this study.

**Figure 4.**
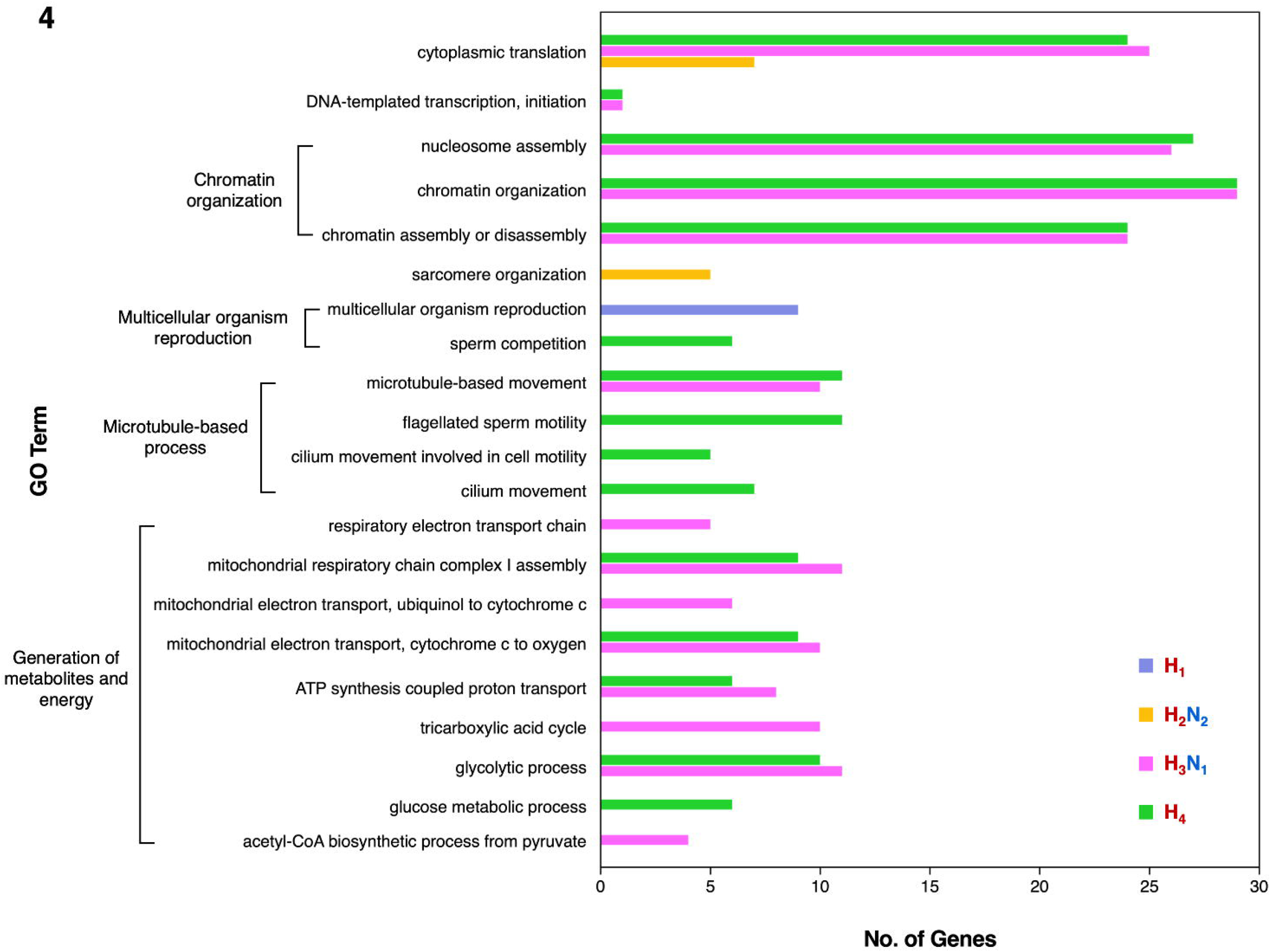
Functional classification of proteins downregulated in response to multigenerational heat stress. Bar graph representing the GOBP terms associated with the proteins downregulated in the stressed sperm proteomes of F_1_ (H_1_) and F_4_ generation (H_2_N_2_, H_3_N_1_, and H_4_), when compared with the unstressed proteome of the F_1_ (N_1_) and F_4_ generation (N_4_), respectively. BP terms with Benjamini-Hochberg corrected p-value < 0.01 and the FDR < 0.01 are shown.

### Effect of multigenerational heat stress on life-history traits

In keeping with our findings that heat stress results in substantial proteomic changes in the *w^m4H^* sperms, we hypothesized that the basic life-history traits, such as reproductive potential and heat stress tolerance of stressed males and their progenies might be impacted. To this end, we first investigated if the fecundity and fertility of the unstressed *w^m4H^* females are affected upon mating with *w^m4H^* males subjected to multigenerational heat stress (**Figure 1A**). Next, to test whether heat stress memory that is multigenerationally inherited imparts any survival advantage to the subsequent generations, we examined the egg-to-adult viability under heat stress or heat tolerance of the progenies of stressed and unstressed *w^m4H^* males at 0-3 hour embryonic stage. Surprisingly, we did not find any significant difference in the reproductive parameters, such as fecundity and fertility, and heat tolerance at the embryonic stage in response to multigenerational heat stress (**Figure S8**).

## DISCUSSION

Since its conception, the phenomenon of MEI has provided a way to elucidate the heritability of traits observed in a non-Mendelian fashion. Although there has been an enormous development in our understanding of the prevalence, underlying mechanisms, and modes of transmission of such effects, it is not in commensuration with the identification of germline components that contribute to these phenotypes in the subsequent generations, especially in *D. melanogaster*. In the present study, we examined how the sperm protein dynamics of *Drosophila* change when subjected to multigenerational heat stress. We found that heat stress given early during embryonic development only for one generation (H_1_) does not result in any substantial effect on the sperm proteome, both in terms of size and enriched biological functions (**Figure 3** and **4**; **Table S4**, **S8**, and **S9**). This finding resonates with the previous report that one generation of exposure to heat stress only results in an intergenerational effect, not multigenerational (8). However, we observed that successive generations of exposure (H_2_N_2_, H_3_N_1_, and H_4_) resulted in a significant downregulation of proteins, with the most prominent effect seen in the sperms of individuals that were exposed to three or more generations of heat stress (H_3_N_1_ and H_4_) (**Figure 3** and **4**; **Table S5-S9**). We speculate that the lower impact of heat stress observed on the sperm proteome of H_2_N_2_ individuals, as opposed to H_3_N_1_ and H_4_, is an outcome of being stress-free for two generations. It is possible that the heat stress-induced epigenetic changes are unstable and slowly return to a default state upon removal of stress, as seen in the eye pigment levels (treatment series #2, **Figure 1B**) in this study and earlier (8).

Sperm cells have long been thought to be translationally silent. However, with advancements in proteomics approaches, sperms in various organisms such as *Drosophila* (46, 47) and humans (48) are now well-established to contain numerous ribosomal proteins (RPs). In *Drosophila*, for instance, 83 RPs have been identified in DmSP3 (46) and 72 by McCullough *et al*. (47) out of 169 annotated so far (Flybase.org). In our study, proteins involved in cytoplasmic translation were one of the significant categories downregulated in response to multigenerational heat stress (**Figure 4** and **Table S9**). We identified only 45 RPs in N_1_ and 43 in H_1_ in the F_1_ generation, perhaps due to a different methodology used in this study. In the F_4_ generation, however, the number was drastically reduced from 34 in N_4_ and 32 in H_2_N_2_ to 9 in H_3_N_1_ and 12 in H_4_, suggesting that the decline in RPs in sperms is due to multiple generations of heat stress at the early embryonic stage, not an artifact. Inhibition and delayed recovery of RP synthesis upon heat stress has been reported previously in *Drosophila* cell line (49). RPs and ribosomal RNAs produce ribosomes, which are essential in maintaining cellular protein homeostasis or proteostasis. Unlike somatic cells, minimal transcription happens in the post-meiotic stages during *Drosophila* spermatogenesis (50–52). Moreover, the translationally repressed mRNAs of essential genes, such as protamines, produced in primary spermatocytes, are translated only at the later stages (26, 53). Therefore, disruption in the RP synthesis caused due to heat stress at the embryonic stage indicates an imbalance in the cellular proteostasis in sperms and possibly elicits a proteotoxic stress response, as seen in the *Drosophila* model of human ribosomopathies (54), budding yeast (55, 56) or mouse (57). Nevertheless, the role of RPs in sperms, post-ejaculation, or post-fertilization remains unclear, and the possibility of their “extraribosomal” role (58) cannot be eliminated without further investigation.

Metabolomics (59) and gene expression (60) studies in *Drosophila* have substantiated that elevated temperature results in alterations in the metabolites involved in energy metabolism, such as glycogen and glucose, as well as the enzymatic machinery. In addition, some perturbations were observed to persist long after alleviation from heat stress. Along similar lines, and with an expectation that metabolism, in general, might be affected by stress, we observed downregulation of proteins involved in glucose homeostasis, tricarboxylic acid cycle, and acetyl-CoA synthetic process from pyruvate in H_3_N_1_ and H_4_ (**Figure 4** and **Table S9**). Recently, a similar decline in the expression of genes involved in metabolism in the offspring of stressed individuals was reported using transcriptome and metabolome analysis in a paternal restraint stress model of *Drosophila* (61), supporting our finding.

The effect of thermal stress and thermal stress-induced oxidative stress on mitochondrial homeostasis has been demonstrated previously in other organisms. It manifests as disturbances in the structure and function of respiratory chain components (62, 63). Due to multigenerational heat stress, components of the mitochondrial respiratory chain and ATP synthase complex are significantly downregulated in sperms in our data (**Figure 4** and **Table S9**). Sperm mitochondria, other than being the powerhouse of cells, undergo major morphological changes during spermiogenesis. Namely, they fuse to form a giant onion-shaped structure called nebenkern and elongate to provide a structural platform on which cytoplasmic microtubules reorganize and facilitate sperm tail elongation (45, 64). Alluding to this, we also found that proteins associated with microtubule-based processes, such as the components of the dynein complex implicated in proper sperm motility and function, are downregulated. Despite this, the fertility potential of the stressed males was not affected (**Figure S8**).

Considering that we observe such changes in sperms, which are temporally separated from the stage of exposure to heat stress (0-3 hour embryo), our results imply an incomplete recovery from the damage caused by repeated environmental insults and point toward the long-lasting effects of MEI. In this regard, our data could not ascribe any adaptive or maladaptive consequences of the MEI of heat stress memory to the subsequent generations (**Figure S8**). Due to the commonality in the gene expression changes associated with aging (65) and proteomic changes observed in this study, they may be manifested as perturbations in other life-history traits, such as lifespan. Therefore, a detailed examination of life-history traits at various stages of the life cycle is needed to understand the evolutionary relevance of MEI.

During spermatogenesis, chromatin is extensively remodeled (45, 66). The crucial step is the replacement of histones by protamines in the late stages of spermiogenesis (53, 67–69). It is well-established that 1-10% and 10-15% of the mouse and human chromatin, respectively, is packaged by histones (70, 71). The degree of histone retention, however, is still unclear in *Drosophila* sperm chromatin. Though undetectable by immunofluorescence (29), there are indications of the presence of histones in *Drosophila* sperms from proteomics and chromatin immunoprecipitation-on-chip (ChIP-on-Chip) studies (46, 72). Notably, we observed downregulation of histone proteins in H_3_N_1_ and H_4_, mainly canonical histone H2A and histone H2B, histone 3 variant H3.3, and histone 4 replacement H4r (**Figure 4** and **Table S9**), which were associated with GO terms ‘DNA-templated transcription-initiation’ and ‘chromatin organization’. Chromatin-based multigenerational epigenetic effects mediated by post-translational modifications of histones have been shown in several studies (14, 17). Although the present study does not directly explore the chromatin landscape of *Drosophila* sperms, the downregulation of histones upon heat stress raises the possibility of loss of histones marks at relevant genomic loci.

For instance, loss of H4r in a null mutant of *Histone 4 replacement* (*H4r*) gene (73) has been shown to act as a mild suppressor of PEV in *Drosophila* and results in more robust induction of heat shock protein (HSP) genes upon heat stress by fine-tuning the chromatin accessibility of heat shock elements and thus providing increased heat-stress tolerance (74). Furthermore, ChIP-seq analysis of H4r in *Drosophila* has revealed its association with regulatory regions of stress response genes such as HSP genes (75), a feature shared with H3.3 (76). Additionally, previous studies in mouse models have reported that nucleosomes associated with sperm chromatin mainly comprise H3.3 (70) and that H3.3 contributes to heterochromatin formation during early embryogenesis (77, 78). Therefore, we speculate that heat stress-induced alterations in histone proteins, presumably chromatin-associated, inherited from sperm to zygote might affect the heterochromatin formation during embryogenesis and, thereby, the expression status of the *white* locus scrutinized in this study.

Apart from histones, we also observed downregulation of other proteins which were associated with the GO term ‘chromatin organization’, such as Protamine B, which packages the paternal genome in sperms (26), versager, which is suggested to colocalize with chromatin during the histone-protamine transition (Flybase.org) and has been shown to affect male fertility in *Drosophila* (79) and Lamin C (80), known to be involved in the regulation of gene expression. Collectively, these findings suggest a role of chromatin-based mechanisms in the MEI of heat stress memory in *Drosophila* via the male germline. While we have been able to capture certain chromatin-associated nuclear proteins, which are affected by heat stress, the sperm nuclear proteome is expected to be much more complex (81), and only improved proteomics approaches (46) or establishing nuclei purification protocols can provide a more detailed molecular insight.

In conclusion, we show that multigenerational heat stress leads to substantial changes in the sperm proteome of *Drosophila*. Our data suggests potential molecular signals and hints toward mechanisms for transmitting the heat stress memory. However, a more thorough analysis is needed by combining the powerful genetic tools of *Drosophila* and advances in omics techniques to truly discern how faithful re-establishment of heat stress memory happens in subsequent generations. Though our proteomics study presents a way forward in understanding the molecular basis of heat stress-induced epigenetic effects in *Drosophila*, MEI is envisaged as a multifaceted phenomenon involving not only proteins but also ncRNAs, DNA modifications, and chromatin components. Therefore, an integrated approach is essential to delineate male gamete-mediated epigenetic effects in *Drosophila*.

## ABBREVIATIONS

ACN: acetonitrile
BP: biological process
ChIP-on-Chip: chromatin immunoprecipitation on chip
DAVID: database for visualization and integrated discovery
DmSP: *Drosophila melanogaster* sperm proteome
FDR: false-discovery rate
GO: gene ontology
HSP: heat shock protein
LC-MS/MS: liquid chromatography with tandem mass spectrometry
LFQ: label-free quantification
MEI: multigenerational epigenetic inheritance
ncRNAs: non-coding RNAs
PEV: position-effect variegation
PIC: protease inhibitor cocktail
RP: ribosomal protein
RT: room temperature
TBST: Tris-buffered saline with 0.1% Tween 20

## DATA AVAILABILITY

The mass spectrometry proteomics data have been deposited to the ProteomeXchange Consortium (http://proteomecentral.proteomexchange.org) via the PRIDE partner repository (82) with the dataset identifier PXDxxxxxx.

## SUPPLEMENTAL DATA

This article contains supplemental data.

## ACKNOWLEDGMENTS

We are grateful to Ravina Saini and Sharath Chandra Thota for their help in the isolation of sperms. We thank the staff of the *Drosophila* Laboratory, Proteomics facility, and Advanced Microscopy and Imaging facility at the Centre for Cellular and Molecular Biology for the technical assistance. S.K. thanks the Department of Science and Technology (DST) for the INSPIRE Fellowship.

## AUTHOR CONTRIBUTIONS

S.K., and R.K.M. Conceptualization; S.K. Methodology, Formal analysis, Investigation, Visualization, Writing – Original Draft; S.K., and R.K.M. Writing – Review and Editing; R.K.M. Supervision and Funding acquisition.

